# Melanin enhances metastatic melanoma colonization by inhibiting ferroptosis

**DOI:** 10.1101/2023.04.03.535376

**Authors:** Arwin Groenewoud, Gangyin Zhao, Maria-Chiara Gelmi, Jie Yin, Gerda E. M. Lamers, Remco van Doorn, Rob. M. Verdijk, Martine J. Jager, Felix B. Engel, B. Ewa Snaar-Jagalska

## Abstract

Melanoma associated death is mainly caused by metastatic disease. Increased melanin levels are associated with decreased melanoma patient survival, yet the contribution of melanin to this process is unknown. Here we show that melanin protects circulating melanoma cells from ferroptosis, enhancing their metastatic potential. We observed that melanin levels in patient-derived uveal melanoma cells as well as cutaneous and conjunctival melanoma cell lines correlate with their metastatic potential in zebrafish xenografts. We find strong associations of the melanin biosynthesis gene *TYRP1*, ferroptosis related enzyme *GPX4* and mitochondrial anion channel (*VDAC1*) with reduced melanoma-specific survival in TCGA data of cutaneous melanoma. Modulation of melanin levels significantly impacts melanoma metastatic potential, increasing or decreasing in concordance with melanin levels. Furthermore, melanin depletion significantly sensitized melanoma cells to ferroptosis leading to a decreased metastatic capacity and enhanced efficacy of ferroptosis induction based anti-cancer therapeutic strategies. Collectively, our results reveal that combined inhibition of melanin biosynthetic enzymes and induction of ferroptosis has potential as a treatment strategy of metastatic melanoma.

**Graphical abstract:** 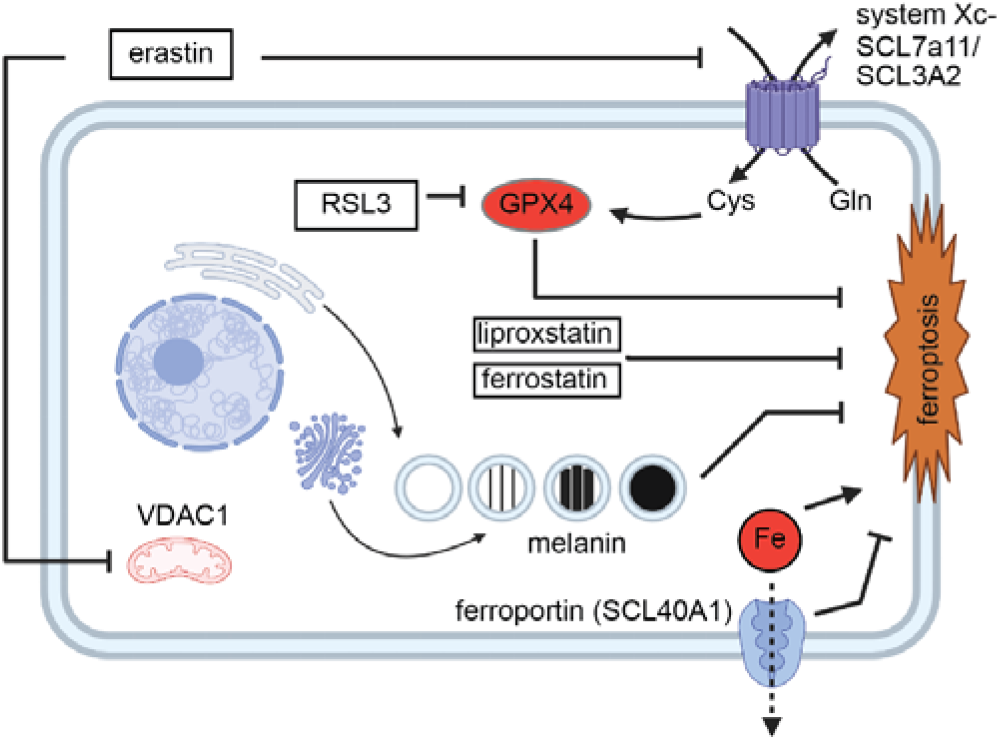

## Introduction

Melanoma is one of the most common malignancies, and arises from melanocytes following malignant transformation. Subsequent metastatic spread, and not the growth of the primary tumor, kills up to 90% of cancer patients^1,2,3^. For (intra-ocular) melanoma, it is assumed that only a small fraction of cells that escape from the primary tumor successfully establish a metastatic colony, indicating a strong selective pressure on disseminating tumor cells within the circulation^4–6^. Among the key factors in curbing metastatic dissemination in the circulation are reactive oxygen species (ROS)^7,8^. These ROS are derived from either intracellular or extracellular stressors, such as but not limited to mitochondrial disfunction, impaired influx/efflux mechanisms, immune or stromal cell interactions^9–12^. In healthy skin, melanin functions to protect against Ultraviolet (UV) radiation by preventing direct DNA damage by absorbing (UV-A/B) thus preventing subsequent ROS-mediated genotoxicity ^13–15^. We hypothesize that melanin protects transformed melanocytes from ROS in a similar manner during metastasis, thus enhancing cell survival during dissemination and the metastatic potential of melanoma cells.

Most melanoma cell lines derived from pigmented lesions lose their capacity to synthesize melanin *in vitro*. Conversely, patient-derived xenograft models retain their melanogenic potential, underscoring the apparent selective pressure *in vivo*. Interestingly, studies suggest an inverse correlation between pigmentation and migratory capacity^21,22^. Strikingly, the *in vivo* cutaneous melanoma model described by Pinner et al. shows an enhancement of distant metastasis in the presence of heightened melanin levels^22^.

Melanomas are derived from neuroectodermal progenitor cells during embryonic development, giving rise to different populations of melanocytic precursors^16,17^. All melanocytes harbor intrinsic melanogenic potential, which is normally induced in the skin in a UV-dependent manner through the αMSH-MITF-TYR axis^16,18^. In ocular melanocytes, melanin biosynthesis is induced through a largely unknown mechanism^18–20^.

One of the possible mechanisms of ROS-mediated cell death is ferroptosis, a lipid peroxidation-based, iron-dependent mechanism of cell death^23–26^. Ferroptosis seems to be more strongly induced in cutaneous melanoma (CM) cells expressing oncogenic RAS variants, possibly due to an increase in cellular iron levels^23,26,27^.

Glutathione peroxidase 4 (*GPX4*) functions as a lipid peroxide reducer, effectively reverting the damage done by ferroptosis^24,25^. Intracellular glutathione is used as a reservoir of ROS reduction and can effectively curb ferroptotic cell death. Common inducers of ferroptosis either interfere with mitochondrial or electron transport chain functions, the cellular system Xc- (erastin), or inhibit GPX4^30^.

During ferroptosis, either induction of mitochondrial stress, endoplasmic reticulum stress through inhibition of voltage-gated anion channel, or inhibition of the cystine/glutamate antiporter system (System Xc-) cause a dramatic increase of intracellular ROS. This sharp increase in ROS catalyzes lipid peroxidation and is presumed to lead to subsequent cell membrane permeation and cell death, while maintaining nuclear integrity^28,29^.

We observed that increased levels of melanin and TYRP1 expression correlate with worse disease outcome in primary UM patients. In addition, melanin inclusion in primary UM tissues increased their respective engraftment capacity in a zebrafish model therefore we propose that the retained melanin protects melanoma cells in the circulation, by scavenging intracellular ROS-mediated damage, preventing ferroptosis, effectively enhancing the metastatic potential of melanoma cells.

To address this hypothesis, we have tested a set of different melanoma cell lines from CM and conjunctival melanoma (CoM) origin with pigmented and non-pigmented phenotypes. We used a melanin depletion strategy for CM and CoM cell lines. Conversely, we developed a biological melanin transfer system, allowing us to re-introduce melanin from a pigmented CM donor cell line to non-pigmented uveal melanoma (UM) cell lines.

We correlated melanin inclusion within UM, CM, and CoM with an enhanced metastatic potential and proved that depletion of melanin in pigmented melanoma cells decreases their metastatic potential. Furthermore, we showed that transfer of exogeneous melanin into UM cells confers protection to stress during hematogenous dissemination. Finally, we demonstrated that melanin depletion significantly enhances susceptibility to ferroptotic insult during dissemination *in vivo*. In conclusion, we confirmed that melanin can act as a pro-metastatic factor protecting cells from ROS and demonstrated the importance of melanin in blocking ferroptosis during metastatic dissemination.

## Results

### The presence of melanin and the upregulation of TYRP1 correlate with tumorigenic potential of UM PDX samples and with a decrease in disease-free survival in UM patients

When studying the metastatic colonization capacity of uveal melanoma in a zebrafish model, we noticed striking differences between tumor samples (Figure 1A). We had generated spheroid cultures from primary UM, as recently described by Groenewoud et al 2023. After dissociation of spheroids, cells were stained with a red transient dye and engrafted intravenously in blood vessel reporter zebrafish larvae (Tg*(fli:eGFP)*) at 48 hours post fertilization (hpf). We measured the engraftment over time based on the fluorescent intensity and the size of the metastatic foci, within the engrafted zebrafish larvae at 1-, 4-, and 6-days post injection (dpi) (Figure 1B). Of the three engrafted PDX samples, the highly-pigmented sample spUm-LB046 showed significant (p<0.001) enhancement of tumor cell number (as measured by fluorescence integrated density) over time, with many tumor cells visible all over the body after 6 days. Tumor sample spUm-LB048 (containing only medium levels of melanin) induced a significant enhancement of fluorescent signal between 1 dpi and 4 dpi (p<0.001), with almost complete tumor clearance at 6 dpi (p<0.001), while the non-pigmented primary UM sample spUM-LB049 was completely cleared from the engrafted zebrafish host at 4 dpi. We considered the option that the differences were caused by different degrees of melanin. As UM tend to spread hematogenously, we verified our findings by assessing the relation between tumor pigmentation and patient survival in a series of enucleated UM (LUMC cohort n=64) (Figure 1C/A) by analyzing the transcription of the terminal enzymatic stages of melanin biosynthesis (Leiden cohort, Figure 1D). Pigmentation levels assessed after enucleation of primary UM sub-divided the tumors into two groups: non- and lightly-pigmented versus medium- and highly-pigmented tumors. Survival analysis indicated that there is a significant increase in melanoma-related death in patients with medium- and highly-pigmented tumors compared to those with non- and lightly-pigmented UM (p=0.006). When comparing melanin biosynthetic genes with melanoma-related death, only the final biosynthetic step of melanin synthesis demonstrated a correlation with bad disease outcome (*TYRP1*, p=0.01) whereas both upstream tyrosinase (*TYR*) and dopachrome tautomerase (*DCT*) were not related to melanoma-related death (*TYR*, p=0.52; *DCT*, p=0.15).

**Figure 1.**
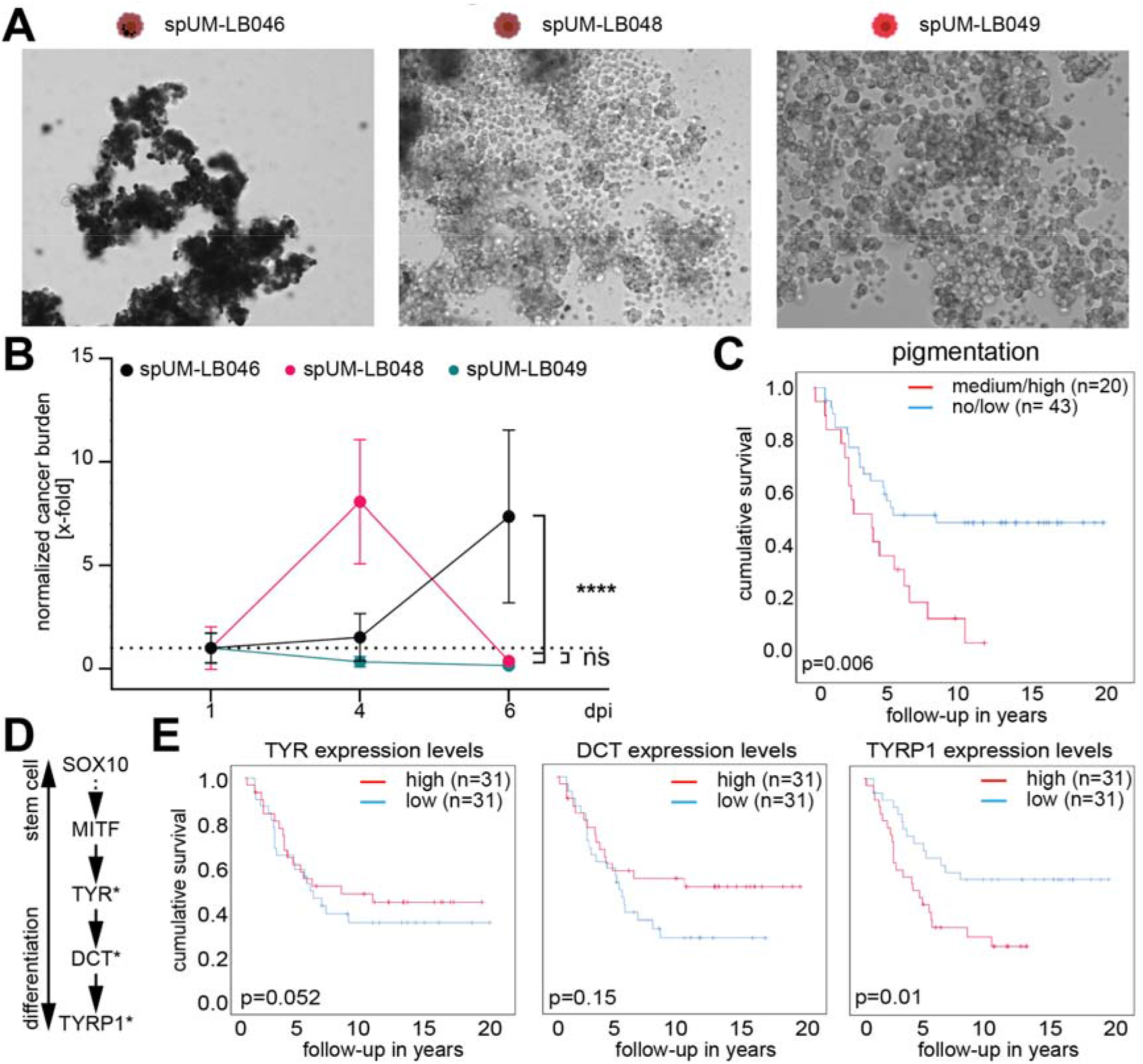
Melanin levels within primary UM cells correlate with survival in vivo. **A)** Three primary UM patient samples ranging from strongly pigmented (spUm-LB046), intermediately pigmented (spUm-LB048) and non-pigmented (spUm-LB049). Melanin levels were derived from phase contrast images of spheroid cultures established from patient material, prior to engraftment. **B)** Three UM spheroid cultures were stained red fluorescent (CMDiI) before intravenous injection into zebrafish larvae and monitored for cancer cell engraftment on days 1, 4, and 6- post injection (dpi). **C)** UM tumor pigmentation at the time of enucleation (histologically determined) and its correlation with UM specific survival. Survival of patients with not and lightly-pigmented tumors (n= 43) was compared to survival in patients (n=20) medium- and highly-pigmented tumors based on pathological assessment. **D)** general differentiation markers used to assess melanocyte differentiation from stem cell to differentiated melanocyte, markers found to significantly affect UM related mortality were marked with an asterisk. **E)** Assessment of the effect of individual melanin biosynthetic genes on UM survival. Expression of the most upstream located tyrosinase (TYR) and the downstream biosynthetic proteins dopachrome tautomerase (DCT, or alternatively TYRP2) and tyrosinase-related protein 1 (TYRP1), analyzed in a group of 64 patients; groups were determined according to the median mRNA expression only TYRP1 expression levels show a negative correlation with survival.

Next, we verified those finding by studying the metastatic colonization capacity of patient derived UM using zebrafish xenografts (Figure 1A). We had generated spheroid cultures from primary UM, as recently described by Groenewoud et al 2023. After dissociation of spheroids, cells were stained with a red transient dye and engrafted in blood vessel reporter zebrafish larvae (Tg*(fli:eGFP)*) at 48 hours post fertilization (hpf). We measured the engraftment over time based on the fluorescent intensity and the size of the metastatic foci, within the engrafted zebrafish larvae at 1-, 4-, and 6-days post injection (dpi) (Figure 1B). Of the three engrafted PDX samples, the highly-pigmented sample spUm-LB046 showed significant (p<0.001) enhancement of tumor cell number (as measured by fluorescence integrated density) over time, with many tumor cells visible all over the body after 6 days. Tumor sample spUm-LB048 (containing only medium levels of melanin) induced a significant enhancement of fluorescent signal between 1 dpi and 4 dpi (p<0.001), with almost complete tumor clearance at 6 dpi (p<0.001), while the non-pigmented primary UM sample spUM-LB049 was completely cleared from the engrafted zebrafish host at 4 dpi. We considered the option that the differences were caused by different degrees of melanin.

Combined these results suggest a causal link between the level of melanin in UM cells and their metastatic potential.

### Ferroptosis resistance marker expression coincides with the expression of melanin biosynthetic enzymes

To assess if our findings in UM, namely that the expression of melanin biosynthetic enzymes correlate with a significant decrease in melanoma specific survival, would be also be applicable and specific to CM and CoM we analyzed available TCGA datasets (Figure 2A, B). We asked if both elevations in expression levels of melanin biosynthetic enzymes or in ferroptosis resistance marker levels showed any significant effect on CM and survival data (Figure 2C, D). We determined the effect of melanin biosynthetic genes on overall survival (Figure 2C). We subsequently focused on the effect of melanin biosynthetic genes on overall survival of CM patients in a group of 458 patients and were able to demonstrate a strong effect of melanin biosynthetic molecules. All melanin biosynthetic pathway genes assessed in this manner in CM patient data, correlated negatively with overall melanoma specific survival (*MITF*, p=0.022; *TYR*, p=0.03; *DCT*, p=0.038; *TYRP1* p=0.0019).

**Figure 2.**
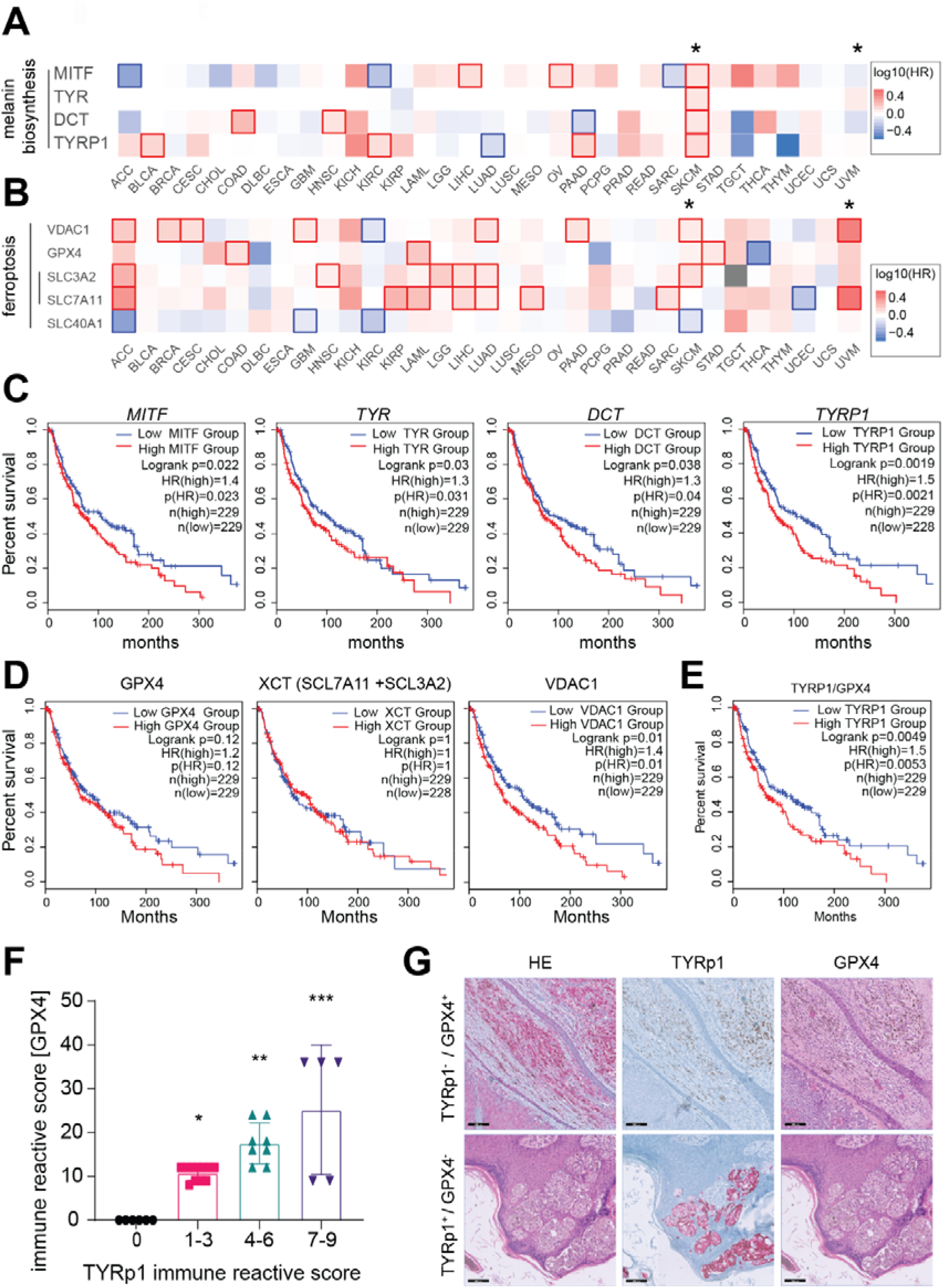
Analysis of melanin biosynthesis and ferroptosis levels in CM melanoma patient. **A)** Survival map of all available TCGA cancer data sets, comparing the effect of melanin biosynthesis genes on cancer associated survival (CM = SKCM and UM= UVM) are marked with an asterisk. TCGA SKCM cohort size = 458, all values are split along the median for each gene. Significant effects on survival are denoted with a red bounding box. In CM melanoma (n= 458), a significant negative correlation with disease free survival for all known melanin biosynthetic genes can be noted (MITF, TYR, DCT, TYRP1). **B)** Comparative analysis of the TCGA, plotting the correlation of disease-free survival for the known anti-ferroptotic mechanisms, system Xc- (SCL7A11 and SCL3A2), GPX4 and mitochondrial VDAC1 show that there is a significant effect of SCL3A2 on survival in CM (p=0.00022) and SCL7A11 in UM (p=0.00014). Data sets are split on the mean and significant effects on survival are denoted with a red bounding box. **C)** Comparative analysis of the effect of melanin biosynthetic gene expression on disease free survival in CM melanoma patients. All known major melanin biosynthetic genes negatively correlate with overall survival (MITF, p=0.022; TYR, p=0.03; DCT, p=0.038; TYRP1 p=0.0019). **D)** Analysis of ferroptosis detoxifying enzymes GPX4 and XCT (SLC7A11 + SLC3A2) and VDAC1 shows that VDAC1 correlates negatively with overall survival. **E)** TYRP1 levels strongly correlate with worse melanoma specific survival when normalized for GPX4 levels. TCGA data, TYRP1 normalized on GPX4 expression, population split along the median. **E)** Analysis of IHC immune reactive scores of TYRP1 on 25 patient tumors stratified on TYRP1 staining levels, enhanced TYRP1 levels correlate significantly with an enhancement in GPX4 staining. Staining intensity * absent = 0, low =1-3, intermediate = 4-6 and high = 7-9. **G)** IHC analysis of human CM patient material, H&E, TYRP1 and GPX4, showing representative TYRP1+/GPX4- and TYRP1-/GPX4+ CM tumors, scale bars denote 200 µm.

Ferroptosis defense mechanisms such as GPX4, XCT (SLC7A11 and SLC3A2) VDAC1 are clinically relevant targets for the induction of ferroptosis. We assessed the association between the aforementioned markers and CM patient survival. We noted that VDAC1 levels significantly correlated with a reduced overall survival for CM (*VADC1* p=0.01; Figure 2D). Subsequently we asked if the co-expression of the melanin biosynthetic enzyme TYRP1 and ferroptosis related transcripts (GPX4, VDAC, XCT) would be associated with a decreased probability of CM specific survival (Figure 2E). To establish the clinical significance of concordant expression of melanin biosynthetic enzyme TYRP1 and ferroptosis resistance markers we compared the association between the overall melanoma specific survival of TYRP1 and TYRP1 normalized to either GPX4, VDAC1 or XCT. Strikingly the negative effects of GPX4 on melanoma dependent survival display a stronger statistical probability after normalization to TYRP1 (p=0.0049) if compared to GPX4 alone (p=0.12, Figure 2E). Subsequently we asked if we could show a similar relation between TYRP1 and GPX4 protein levels in patient material, in a selection of different cutaneous melanoma IHC samples (derived from patients with different skin types, n=30) we assessed immune reactive scores (staining intensity multiplied by staining percentage, binned in absent TYRP1 (0), low (1-3), medium (4-6) and high (7-9)) of both markers and were able to find a significant correlation between TYRP1 and GPX4 expression (presence of GPX4 staining in TYRP1 absent compared to low, p=0.0124; absent compared to medium, p=0.001 and absent compared to high p<0.001) (Figure 2F, G).

### Intracellular melanin and TYRP1 levels predict cancer cell engraftment

To further test our hypothesis that the presence of intracellular melanin plays a role in metastatic dissemination of different types of melanomas, we analyzed a matched panel of pigmented and non-pigmented melanoma cell lines (schematic representation in Figure 3A). The metastatic colonization in zebrafish was measured at 1, 4 and 6 dpi, comparing pigmented (marked by an *) and non-pigmented cell lines derived from CoM (CRMM1*, CRMM2), CM (SK-Mel28, PDX11917*) and UM (XMM66 and OMM2.3). All tested cell lines were transduced with lentiviral tdTomato and data was analyzed after normalization of fluorescent intensity to 1 dpi (as described previously)^33^. For both CoM and CM, we noted a significantly enhanced metastatic colonization for the pigmented cell line when compared to the non/low pigmented cell line within the cell line pairs (figure 3B, C). Metastatic colonization was not exclusively linked to melanin content, CRMM1*, p<0.001 when compared to CRMM2. Conversely SK-Mel28, greatly showed enhanced metastatic capacity p<0.001 when compared to CM patient derived xenograft culture PDX11917. Cell line PDX11917 was found to be overtly pigmented, both in culture and when concentrated during sub-cultivation, showed marginal proliferation in zebrafish, and was able to establish metastatic colonies at 6 dpi^34^. UM lines that were used in this panel were deemed to be non-pigmented and as expected failed to form any metastatic colonies as reported by Groenewoud et al 2021. Subsequently, we determined if there were detectable melanosomes within the cell lines that made up our panel, to ensure that all our designated non-pigmented cell lines indeed did not contain any melanosomes (Figure 3D). Strikingly, not only CRMM1 and PDX11917 but also SK-Mel28 showed active melanosome formation when observed using transmission electron microscopy (TEM). In line with our previous findings, we asked if in this cell line panel, we could correlate *TYRP1* mRNA levels with their metastatic capacity (Figure 3E). We noted a significant increase of *TYRP1* expression in CRMM1, PDX11917 and SK-Mel28 when compared to their non-pigmented counter parts or when compared to the non-pigmented cells within this panel (p<0.001). None of the non-pigmented cell lines (CRMM2, XMM66, OMM2.3) showed detectable *TYRP1* mRNA, indicating that both TEM and qPCR should be used to verify the presence or absence of melanogenic capacity of melanoma cells.

**Figure 3.**
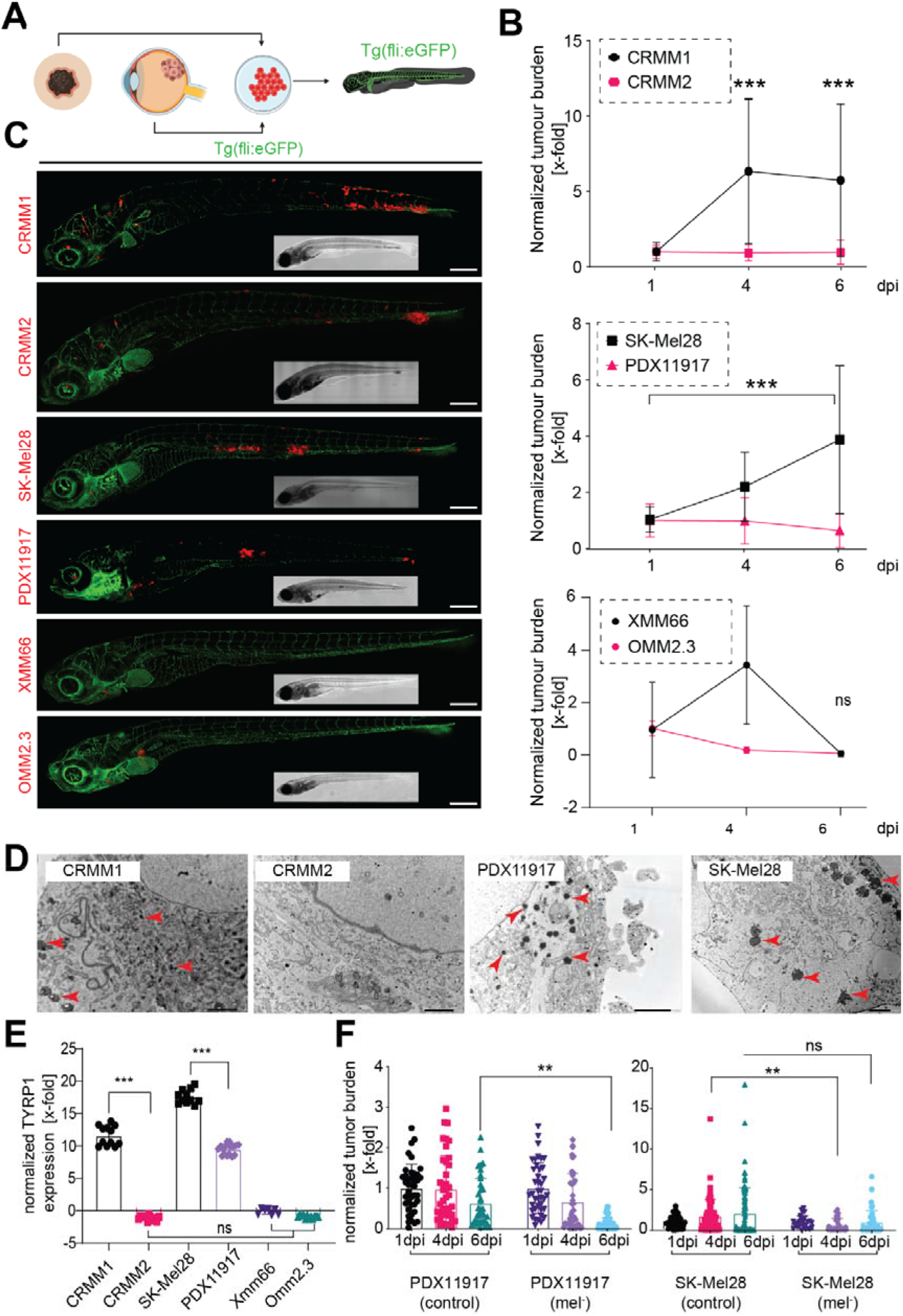
Engraftment of a (pan)melanoma panel in Tg*(fli:eGFP) x Casper* zebrafish shows efficient engraftment of CM and CoM melanoma, whereas UM is readily cleared from zebrafish. **A)** Schematic representation of the experimental approach: CoM, CM and UM cell lines were injected in to Tg*(fli:eGFP)* blood vessel reporter zebrafish through the duct of Cuvier at 48 hours post fertilization. Pairs of pigmented and non-pigmented cell lines were chosen for CoM and CM and two non-pigmented cell lines for UM. **B)** Growth kinetics of the (pan)melanoma cell line panel, after hematogenous engraftment into zebrafish. For each experiment, 40 individuals were imaged, divided over two biological replicates. For all measurement, the integrated fluorescence density was plotted for 1, 4 and 6- dpi (all measurements were normalized to day 1 of the individual cell line). Measurements shown are the mean, error bars represent ± SEM. **C)** Confocal micrographs of representative phenotypes of the engrafted cell lines at 6 dpi, scale bar is 250 µm **D)** Transmission electron micrographs of CoM and CM melanoma cell line pairs; (type IV melanosomes, indicated with ▴). Scale bars are 2 µm, all images are representative images. **E)** Quantitative PCR measurements of tyrosine related protein 1 (TYRP1), the enzyme responsible for the terminal biosynthetic conversion of tyrosine into melanin. **F)** Melanin depletion through PTU treatment of pigmented and non-pigmented cells. CM melanoma PDX-derived cell line PDX11917 and non-pigmented melanoma cell line SK-Mel28 were depleted for 14 days prior to engraftment through injection into zebrafish. The mean and the standard error of the mean (SEM) were plotted, n=20*2. p<0.05=* p<0.01=** p<0.001=***.

Subsequently, we measured the effect of chemical inhibition of melanin biosynthesis on the metastatic colonization of highly-pigmented PDX-derived CM cell line PDX11917 when compared to SK-Mel28, a cell line bearing only occult melanin. We demonstrated that PDX11917’s metastatic capacity was significantly inhibited after chemical melanin depletion, by treatment with N-Phenylthiourea (PTU) contrary to vehicle control, at 6 dpi (p<0.01). SK-Mel28 was not significantly inhibited by melanin depletion at 6 dpi, but displayed a significant delay in metastatic colonization at 4 dpi (p<0.01) which recovered by 6 dpi, and indicated no overall inhibition when compared to untreated control (Figure 3F).

These observations suggest that the presence of melanin enhances a melanoma metastatic capacity and that TYRP1 levels are indicative of melanin biosynthesis, given that upstream activation is present. Strikingly, SK-Mel28 shows strong expression of TYRP1, but has only minimal melanosome formation and occult melanin, as visualized under TEM, under normal culture conditions. This further implies that the presence of melanin rather than solely the expression of TYRP1 is required for the enhancement of tumorigenic capacity.

### Melanin depletion of highly-pigmented CM melanoma cell line mugmel2 significantly reduces its metastatic potential

After the primary assessment of the effect of melanin on metastatic dissemination of melanoma cells in zebrafish xenografts, we asked if this phenomenon could be validated in a highly-pigmented *NRAS-*mutated CM cell line: mugmel2^31^. We first determined the presence of all stages of melanosome maturation in mugmel2 cells, using transmission electron microscopy (TEM) as shown in (Figure 4A). Melanosomes Type II, III and IV were readily visualized due to their intrinsic electron density, and are indicated with yellow (▲ type II), blue (▲ type III) and red (▴ type IV) arrowheads, respectively. The eGFP-expressing mugmel2 cells were engrafted in zebrafish as described previously, with or without prior chemical depletion of melanin through 1-phenyl 2-thiourea (PTU). Intravenous injection of the melanin-depleted mugmel2 cells induced less metastatic colonization in *Casper* zebrafish at 6 dpi (Figure 4B, C) in comparison to non-depleted cells *in vitro* (Figure 4C). Next, we measured the decrease of metastatic potential at 4 dpi and 6 dpi for two concentrations of PTU (125 and 500µM) when compared to treatment with equivalent volumes of vehicle control (DMSO).

**Figure 4.**
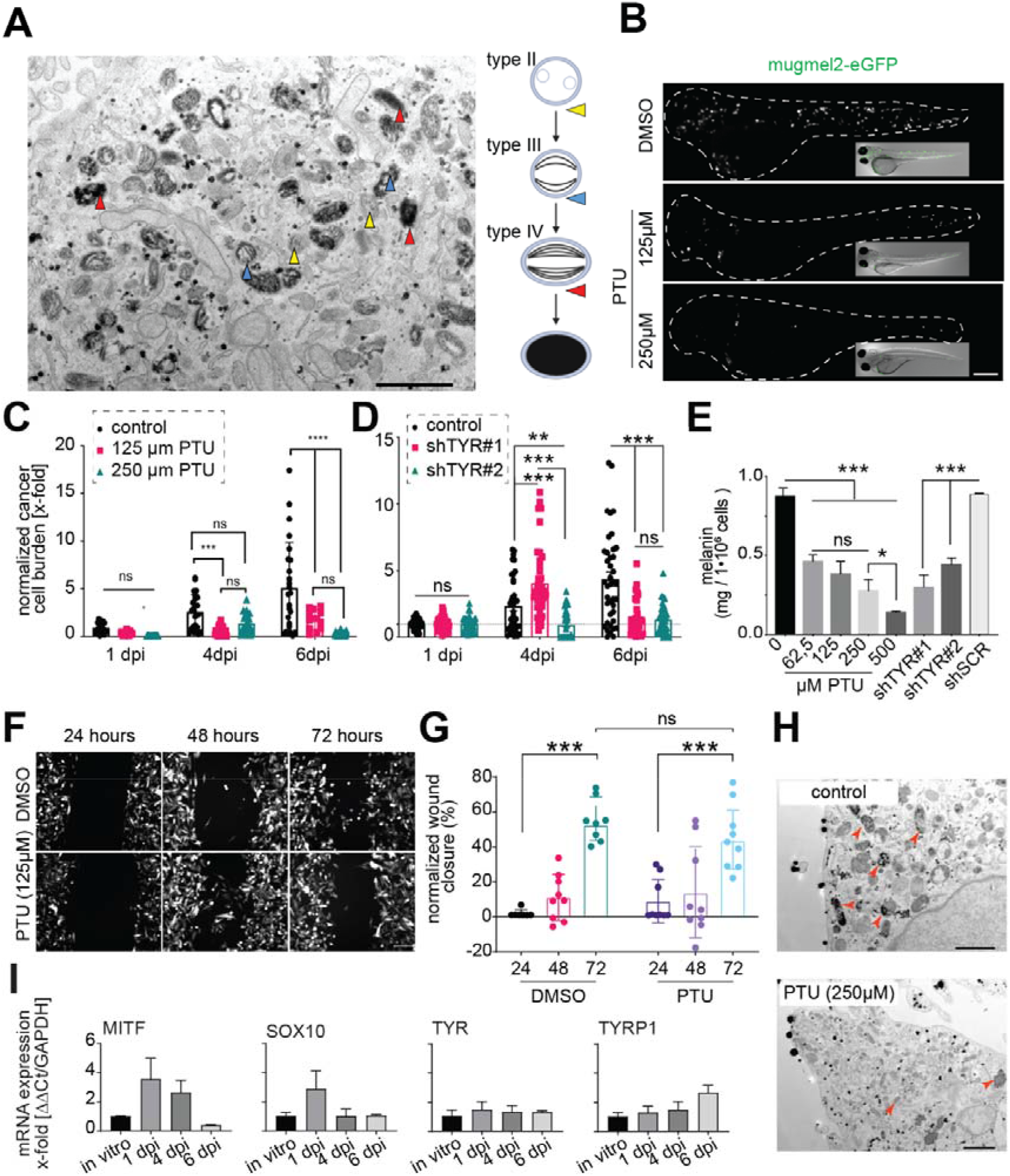
Melanin depletion of pigmented melanoma cell lines decreases their tumorigenic potential. **A)** Transmission electron microscope (TEM) assessment of melanosome maturation in mugmel2 cells: type II melanosomes are indicated by ▴, type III melanosomes by ▴ and type IV by ▴. Scale bars are 2 µm, and all images are representative images. **B)** Engraftment of melanotic melanoma cell line mugmel2, labelled lentivirally with eGFP, in non-pigmented *Casper* zebrafish, representative confocal micrographs were shown. Mugmel2 cells were treated with 1-phenyl 2-thiourea (PTU) in vitro prior to engraftment and its effect on cell intrinsic metastatic potential was compared to DMSO control **C)** PTU inhibition of mugmel2 melanation and its effect on the metastatic capacity of mugmel2 in vivo. Cells were depleted in vitro through PTU addition 2 weeks prior to hematogenous engraftment into Casper zebrafish (n=2 * 20). Measurements were normalized to 1 day post injection (dpi), engraftment was monitored on 1, 4 and 6- dpi. **D)** Quantification of cancer cell engraftment of zebrafish implanted with mugmel2-eGFP, containing shSCR, shTYR1#1 or shTYR1#2. Measurements were normalized to day 1 of each individual condition. **E)** Dose-dependent melanin depletion upon in vitro application of PTU to mugmel2 cells compared to solvent control and genetic depletion of TYR (lentiviral shRNA mediated knock down) compared to scrambled short hairpin control, as measured by spectrophotometer. **F)** Cellular migration (wound healing) of mugmel2 cells treated with solvent control (DMSO) compared to PTU-mediated chemical depletion of melanin, shown as representative epifluorescent micrographs and in panel **G)** as quantification of wound area over time, normalized to wound area at t=0. **H)** TEM micrographs noting the chemical depletion of melanin and the subsequent reduction of visible melanosomes when compared to solvent control. **I)** *ex vivo* qPCR quantification of melanocyte differentiation markers MITF, SOX10, TYR and TYRP1, on engrafted mugmel2 cells following FACS isolation from metastatic colonies. Respective quantifications are normalized to transcription levels *in vitro* and were normalized internally to GAPDH levels (were normalized internally to GAPDH levels and were normalized internally to GAPDH levels (ΔΔCt). All measurements were generated from 3 biological replicates (pooled from 100-300 larvae prior to FACS isolation). The mean and the standard error of the mean were plotted (SEM), n≥20*2. Statistical significance was indicated as p<0.05=* p<0.01=** p<0.001=***.

Genetic shRNA mediated interference with melanin biosynthesis through inhibition of tyrosine (*TYR)* by led to a significant inhibition of metastatic capacity when compared to scrambled shRNA control concordantly (Figure 4D, p<0.001).

Subsequently we measured the concentration of intracellular melanin spectrophotometrically. For the chemical depletion the strongest inhibition, without deleterious effects on cell survival, was induced by 500 µM PTU (approximately 85%, p<0.001) and a dose-dependent increase in melanin content was recorded between 500 µM and lower PTU concentrations (p<0.05) (Figure 4E). Both shTYR constructs were sufficient to reduced intracellular melanin levels significantly (Figure 4E)^20,32^.

To assess whether this decrease in metastatic capacity could be due to a decrease in overall migration, we measured cell migration *in vitro* using a wound healing assay. We observed no difference in the migratory potential when comparing PTU-treated cells with the vehicle control (Figure 4F, G). Using TEM, we determined whether PTU treatment induced loss of melanosomal structures in mugmel2 cells (Figure 4H). We observed a loss of type II, III and IV melanosomes upon PTU treatment (250µM). In summary, these data clearly suggest that the degree of melanation of mugmel2 correlates with its metastatic potential. The presence of all known stages of melanosomes indicates that cell line mugmel2 has retained its canonical melanogenic phenotype. To determine if the mugmel2 metastatic capacity was driven by a de-differentiation, or general enhancement of stem cell like features, we assessed mRNA transcription levels *ex situ* (tumor explants of 50 individual tumors per timepoint, n=3). We determined the transcript levels of MITF, SOX10, TYR and TYRP1, compared to *in vitro* cultured mugmel2 cells, to assess any enhancement of stem cell like properties (MITF, SOX10) or any differentiation/melanin biosynthesis markers (TYR, TYRP1) induced by interaction with or selection by the zebrafish microenvironment. We noted that over time stem cell like properties seem to diminish where melanin biosynthesis markers are slightly enhanced by the end of the experiment (6 days). Taken together these data suggest that the depletion of melanin from mugmel2 through either chemical or genetic means significantly decreases its tumorigenic capacity. This depletion does not significantly alter its migratory capacity, nor does it enhance stem cell properties. This implies that not migration, but subsequent steps in metastatic dissemination *in vivo* are affected by the presence of intracellular melanin.

### Introduction of extraneous melanin is sufficient to re-establish UM metastatic potential

We previously observed and described that UM cell lines are generally non-metastatic, non-pigmented and do not express TYRP1. Many UM patients have a dark brown to black tumor at the time of enucleation, and we noticed that both a high level of pigmentation as well as a high TYRP1 expression correlate with poor survival. Furthermore, the tested pigmented primary samples were capable of establishing metastatic colonies in zebrafish (https://doi.org/10.1101/2021.10.26.465874).

To this end we asked if introduction of extraneous melanin would be sufficient to re-instate the, in patients clearly noticeable, metastatic potential UM cells. Therefore, we established a co-culture system to allow *in vitro* transfer of melanin from donor pigmented cells (mugmel2) to naïve UM cells (XMM66, OMM2.3) (Figure 5A, Supplementary Figure 1). We included a co-culture with vehicle (DMSO) in parallel to a melanin-depleted co-culture. Donor cells were pre-incubated with Mitomycin-C, a mitotic spindle poison, abrogating mitotic potential, while retaining overall cell viability; this allowed us to perform a protracted (72 hours) co-culture with the donor cells while blocking donor cell outgrowth. To determine how much melanin would be taken up, we measured the intracellular melanin concentration in naïve UM cultures, UM cells co-cultured with pigmented mugmel2 (mel+) and chemically melanin-depleted mugmel2 (mel-) cells. We noted a significant melanin enhancement in pigmented co-cultures, for both UM cell lines compared to both naïve (2 to 3-fold) and melanin-depleted co-cultures (8 to 10-fold approximately) (Figure 5B). Using TEM, we verified the successful transfer of intracellular melanin from highly-pigmented melanoma cell line mugmel2 (mugmel2 mel+) to XMM66 cells (Figure 5C). In naïve XMM66 cells, melanosomes were completely absent, whereas large melanosomal structures were observed in the co-cultures with pigmented cells. In contrast, in the melanin-depleted co-culture, only a few small electron dense vesicles and some empty vesicles were observed (Figure 5C), indicating functional transfer of melanin from pigmented CM donor cells to naïve non-pigmented UM cells (additional TEM images are presented in supplementary Figure 2). Subsequently, we asked if the transferred intracellular melanin could play a protective role after UM cell injection into the bloodstream of zebrafish. We therefore injected these sets of co-cultured cells Xmm66 and OMM2.3 labeled with red fluorescent, co-cultured with green mugmel2mel+ and mugmel2mel-cells, into zebrafish and scored the metastatic burden at 6 dpi. In pigmented co-cultures, both UM cell lines gained a significant enhancement in metastatic colonization in contrast to the melanin-depleted co-cultures (Figure 5D, E). Importantly these results prove that melanin inclusion into UM cells rescues their survival in circulation leading to metastatic dissemination.

**Figure 5.**
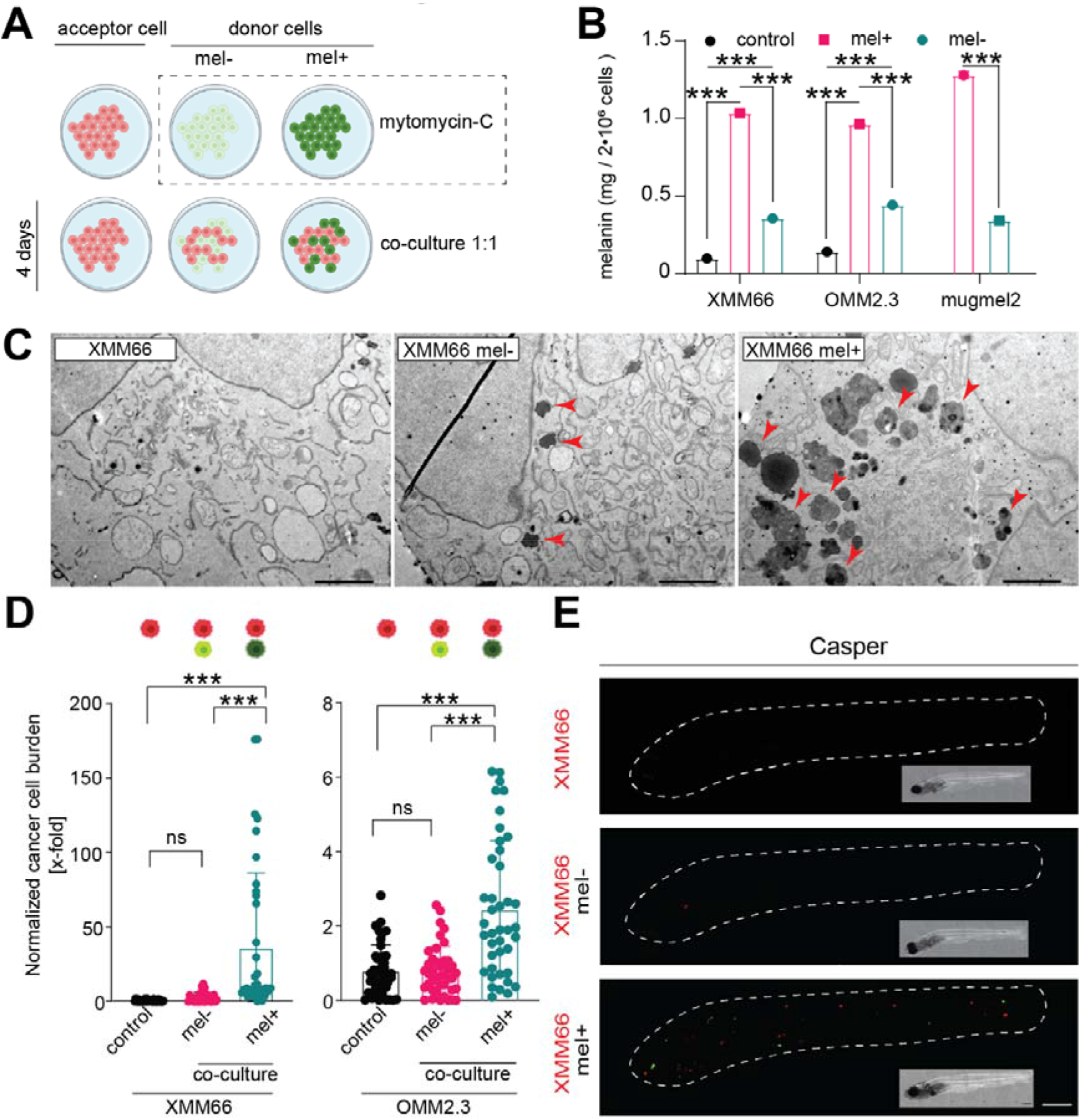
In vitro melanin transfer from donor cells into recipient UM cells rescues their metastatic potential *in vivo*. **A)** Schematic representation of melanin transfer co-culture model. Recipient (red, UM cells) and donor cells (green, mugmel2, mugmel2 mel+, pre-treated with DMSO and mugmel2 mel-melanin depleted through pre-treatment with PTU) were cultured separately. Prior to co-culture, donor cells were pre-treated with mitomycin-C (100µg/mL) for 3 hours. Cells were mixed in a 1:1 ratio of acceptor cell combined with mel+ or mel-mugmel2 cells. After 4 days of co-culture, cells are harvested and either engrafted into zebrafish or used for in vitro analyses. **B)** Spectrophotometric analysis of uptaken melanin in UM cells, calculating mg / 2* 106 cells. 3 biological repeats. **C)** Representative transmission electron micrograph indicates the internalized melanosomes in UM cells donated from mugmel2 mel+ cells (type IV melanosomes, indicated with ▴). Scale bar represents 2 µm. **D)** End point measurement (6 dpi) of zebrafish (n=2 x 20) engrafted with naïve cells (control), UM cells co-cultured with melanin-depleted donor cells (mel-) and co-cultured with pigmented donor cells (mel+). **E)** Representative fluorescent micrographs of co-cultured UM cell line XMM66, show the naïve XMM66 cells (red), XMM66 cells co-cultured with green melanin-depleted (mel-) and melanin-proficient (mel+) mugmel2 donor cells. Some mugmel2 cells (green) remain in circulation, do not form metastatic colonies but increase survival of UM cells as indicated in D.

### Melanin protects against ferroptosis *in vitro* and *in vivo*

Successful metastatic colonization is a relatively rare occurrence in most cancers and can be described as a stochastic process. Here, random chance aligns with the metastatic cells intrinsic properties to allow a minute subset of cancer cells to form a metastatic colony in a suitable niche. Recent discoveries have highlighted the importance of reactive oxygen species (ROS) and more specifically ferroptosis in the curbing of metastatic dissemination in CM^7,35^. Since melanin has long been known to act as a ROS scavenger^18,19^ and we have shown within this manuscript that upregulated melanin biosynthesis correlates with worse melanoma specific disease outcome. Based on this we hypothesized that melanin might scavenge ROS while cells are in the circulation, prolonging circulating tumor cell (CTC) survival, effectively enhancing their chance of finding a suitable niche.

To address this hypothesis, we first tested the effect of melanin depletion on cell membrane peroxidation *in vitro*, as this is one of the hallmarks of ferroptosis. Cell membrane oxidation marker BODIPY 581/591 was used to compare the pro-ferroptotic effects of XCT/VDAC or GPX4 inhibition with or without rescue with membrane peroxidation inhibitor ferrostatin-1 (Figure 6A). In both XCT/VDAC and GPX4 inhibition groups (utilizing erastin or RSL3 respectively) we noted a significant increase of membrane peroxidation in the melanin depleted cells when compared to the melanin proficient groups. In melanin depleted cells the addition of 2µM ferrostatin-1 was sufficient to significantly reduce membrane peroxidation, in melanin proficient cells this reduction was not significant (Figure 6B).

**Figure 6.**
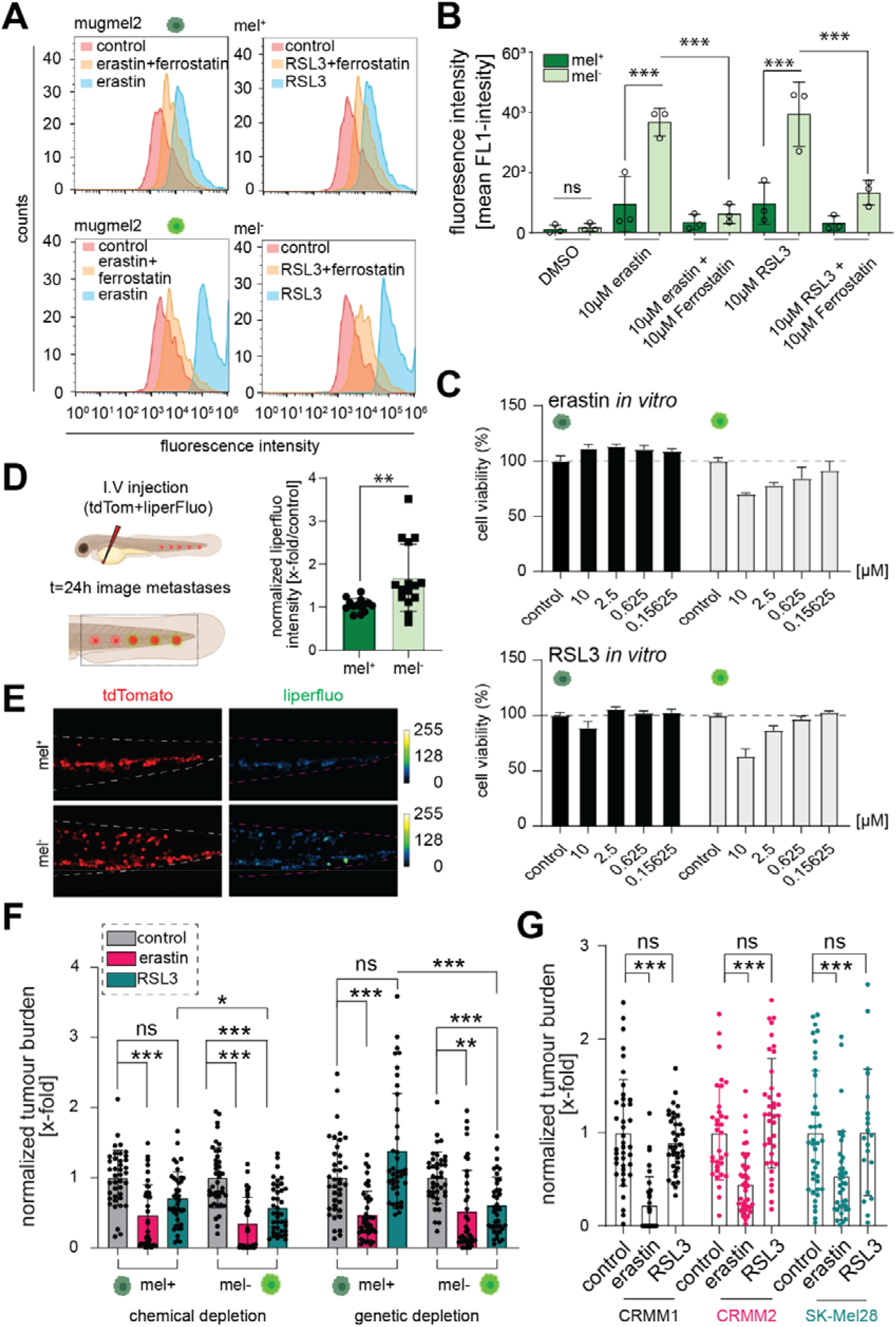
ROS and ferroptosis is quenched by intracellular melanin, erastin induces ferroptosis at sufficient levels to overcome cellular ROS defenses. **A)** Membrane lipid peroxidation measurements using BODIPY 581/591 of mugmel2 cells with (mel-) and without melanin (mel+) through chemical inhibition (PTU pre-treatment), as described. Both cells with and without melanin were treated with Erastin (5 μM), RSL3 (10 μM), Erastin (5 μM) + Ferrostatin-1 (2 μM) or RSL3 (10 μM) + Ferrostatin (2 μM) 8 hours prior to flowcytometric measurement. **B)** Quantification of fluorescent (BODIPY 581/591) signal. **C)** In vitro proliferation (WST1) assay to assess the growth inhibitory effects of ferroptosis inducers erastin and RSL3 on mel+ and mel-mugmel2 cells. **D)** in vivo measurement of lipid peroxidation of circulating mugmel2 cells after engraftment (24h), cells (tdTom^+^) were stained lipid peroxidation reactive dye (Liperfluo, 10µM) for one hour prior to engraftment. **E)** quantification of Liperfluo intensity of tdTom^+^ mugmel2 cells in circulation. Liperfluo signal intensity was measured, using a confocal microscope, in 3 experimental replicates, focusing on cells in the caudal hematopoietic tissue. Signal intensity was rescaled and show as an intensity plot (representative individual, rescaled to 0-255 bits). **F)** *In vivo* ferroptosis induction as described in D in larvae engrafted with naïve mel+ and depleted mel-mugmel2 cells after chemical (PTU) and genetical (shTYR#2) melanin depletion prior-engraftment. Melanin depletion by chemical and genetics means sensitized melanoma cell lines to RSL3 whereas erastin mainly seems to circumvent or overpower ferroptosis resistance mediated by melanin. **G)** Ferroptosis induction in ZF xenograft models *in vivo* obtained by engraftment of a melanoma panel with CoM (CRMM1 and CRMM2) and CM cell lines (SK-Mel28 and mugmel2). Ferroptosis inducers erastin and RSL3 were added to the previously determined maximum tolerated dose (MTD) (results not shown, manuscript in writing, erastin 5 µM and RSL3, 10 µM) to the zebrafish water of engrafted larvae at 3 dpi. The water containing the compounds was exchanged every other day. At 6 dpi, the cancer cell burden was measured and subsequently normalized to the vehicle control group (of each individual condition).

To assess the specific growth inhibitory effect of known ferroptosis-inducers erastin and RSL3, we performed a WST1-based proliferation assay on both melanin proficient and deficient mugmel2 cells (Figure 6C). Strikingly, we noted that under normal culture conditions melanin proficient mugmel2 cells are largely refractory to ferroptosis induction with erastin or RSL3. We found that only melanin-depleted mugmel2 was significantly sensitized to both erastin (approximate growth inhibition 70-80%, 10-0.625 µM, p<0.001) and RSL3 (approximate growth inhibition 60-80%, 10-1,25 µM, p<0.001 and 0.625 at p<0.05). In contrast, the melanin-proficient mugmel2 were only susceptible to the highest concentrations of RSL3 (10 and 5 µM, p<0.001).

To determine if we can measure a similar increase in membrane peroxidation *in vivo* we developed an *in vivo* membrane peroxidation assay (Figure 6D, E). To this end, we stained tdTomato expressing mugmel2 cells (both melanin pro- and deficient) with membrane peroxidation specific dye Liperfluo 1 hour before engraftment. We imaged both engrafted populations using a confocal microscope, focusing on the caudal hematopoietic tissue (CHT), the location where the majority of the metastatic colonies will form. Within 24 hours post engraftment we measured a significant difference between melanin pro- and deficient mugmel2 cells, where the latter showed a significant enhancement when compared to the melanin containing mugmel2 cells.

We engrafted zebrafish larvae with either melanin pro- or deficient mugmel2 cells (depleting melanin through chemical and genetic means) and treated the engrafted larvae for 6 days (1 - 6 dpi) with 5 µM erastin and 10 µM RSL by water administration (Figure 6F). We measured the tumor burden (total red fluorescent objects ≥10um within the zebrafish larvae), normalized to vehicle control (DMSO) at 6 dpi. There was no significant sensitization to erastin, most likely due to its strong inhibitor capacity on both melanin pro- as deficient mugmel2 cells. Strikingly RSL3, acting through inhibition of GPX4, significantly reduced the tumor burden in both chemically and genetically-melanin depleted populations, indicative of resistance to canonical ferroptosis induction mediated by the presence of intracellular melanin.

To determine the cross-sample validity of our hypothesis, we engrafted zebrafish larvae intravenously with our previously described set of melanoma cells (UM, CM and CoM), at 48 hpf and treated the engrafted larvae from 1 dpi with 5 µM erastin and 10 µM RSL (Figure 6G). We subsequently measured the tumor burden at 6 dpi and normalized the tumor burden to the vehicle control (DMSO). We noted that CoM cell lines CRMM1* and CRMM2 and CM cell line SK-Mel28* showed a significant (p<0.001) reduction of tumor burden when treated with erastin. RSL3 did not significantly reduce tumor burden in any of the tested cell lines.

In conclusion, these data suggest that melanin protects melanoma cells in circulation, functionally mitigating intracellular ROS and protecting circulating cancer cells from ferroptotic cell death, indirectly enhancing tumorigenic capacity.

## Discussion

Melanomas are often strongly pigmented in patients. The prevalence of melanoma pigmentation underlines not only the cellular lineage they derive from, but also the presence of selection pressure forcing the expression of melanin-related genes. Cells derived from pigmented melanomas generally lose their melanin biosynthetic capacity and increase their migratory capacity *in vitro*^22^. Next to clinical correlations that associate either melanin or melanin-related gene signatures with an unfavorable disease outcome, the true functional role of melanin in melanoma development remains unclear and contradictory^21,36^.

We engrafted spheroids derived from primary UMs tissues in zebrafish and observed that there was a relation between the presence of melanin in the primary UM samples (clinically scored for melanotic level in a +, +/- and – scale) and prolonged circulation and tumorigenic potential in zebrafish after engraftment. This experiment indicated that there is a significant difference in tumorigenic potential of heavily-pigmented patient samples when compared to both intermediate and lightly-pigmented samples. Interestingly, UM cell lines OMM2.3 and XMM66, originally derived from metastatic UM, had lost all melanogenic capacity *in vitro*, and were readily cleared from the engrafted host after systemic injection (within 16 hours post injection). This observation and the short timeframe, wherein near complete attrition of circulating tumor cells (CTCs) is attained, is in line with a possible induction of ROS-mediated cell death^8^. Strikingly, we observed strong tumorigenic capacity when engrafting low passage spXMM66 cells, derived from a pigmented patient-derived xenograft tissue (PDX), while an immortalized cell line from the same patient XMM66 proved to be non-tumorigenic. In addition, our analysis of a clinical UM cohort confirmed a strong association between high tumor pigmentation, high expression of *TYRP1*, the terminal melanin biosynthetic, and high UM-related death in patients (LUMC cohort n=64).

Following this observation, we reasoned that this phenomenon might hold true for other melanoma sub-types. We concordantly observed that there has been a nearly complete loss of melanin biosynthesis in most melanoma cell lines, presumably through negative selection (or general lack of selection pressure) by successive *in vitro* cultures without selection pressure.

To assess this, we acquired the aberrant CM cell line mugmel2, an NRAS-driven and heavily-pigmented melanoma cell line^31^. We assessed the formation of melanosomes using TEM and measured melanin levels using a spectrophotometric method described by Friedmann et al^18^. Both chemical and genetic inhibition of melanin biosynthesis, by treatment with 1-Phenyl-2-thiourea (PTU) or *TYRP1* shRNA interference, reduced melanin levels and decreased metastatic dissemination in zebrafish xenografts.

We assembled a set of UM, CM and CoM melanoma cell lines and were able to generate a set of paired pigmented and non-pigmented CM and CoM cell lines. We repeated the chemical depletion of melanin in the CoM melanoma set and found that also within this set there is a significant reduction of tumorigenic potential upon depletion of melanin, suggesting that melanin has a canonical pro-tumorigenic function in all melanomas. Strikingly, our findings validate the work from Pinner et al 2009, where they state that low melanation correlates with high migratory capacity of CM cells *in vivo*, but that pigmented cells do generate more distant metastases^22^.

To assess the mechanistic effect of melanin levels on the metastatic behavior of UM cell lines we developed a co-culture system to transfer melanin from mugmel2 donor cells into UM acceptor cells. Through this system, we introduced exogenous melanin to UM cells, for subsequent assessment of metastatic capacity. Other methods to re-instate melanin biosynthesis were unsuccessful (treatment with a-MSH, forskolin or lentiviral over-expression of melanin biosynthetic genes *DCT*, *TYR* and *TYRP1*, results not shown). Spectrophotometric measurement of the uptake of melanin and visualization of melanosomes through TEM confirmed that UM cells readily take up melanin from external donors. Co-cultures of both melanin biosynthesis proficient and deficient donor cells were generated and we measured a significant enhancement of detected melanin in co-culture with melanin proficient donors when compared to melanin deficient donors or untreated control cells. Both OMM2.3 and XMM66 co-cultured with melanin-proficient cells showed a significant enhancement in tumorigenic potential when compared to either co-culture with melanin deficient or untreated cells. These findings indicate that UM cells are capable of up taking melanosomal melanin, at least *in vitro,* and underscore one of the possible functions of melanin in the distant metastasis of UM.

Taking together our observations with recently published experimental proof that circulating melanoma cells are largely killed through reactive oxygen^7,35^, we hypothesized that ROS-based cell death might be the underlying mechanism driving UM attrition in circulation^7^. In line with this hypothesis, melanin inclusion within UM cells might thus help prevent *in vivo* cell death of circulating UM cells. More recently there has been experimental proof that circulating tumor cells are killed specifically by an iron-dependent non-apoptotic cell death mechanism known as ferroptosis^35^.

Using our model, we tested if melanoma cells were responsive to the induction of ferroptosis during their time in circulation in the zebrafish model. We started out by challenging duplicate sets of pigmented and non-pigmented melanoma cells we used previously and demonstrated a clear correlation between the levels of melanation and the response to the induction of ferroptosis with inhibitors of GPX4 ((1S,3R)-RSL3; alternatively named RSL3) and system Xc- (erastin)^23,25,27,37^. Ferroptosis induction by erastin occurs through perturbation of system Xc- and mitochondrial voltage-dependent anion channels^26,37^. The *in vivo* induction of ferroptosis with erastin proved to be highly effective, reducing tumor burden independent of melanin inclusion for cell line SK-Mel28, which showed low level melanation under TEM. Additionally, erastin proved sufficiently potent to induce ferroptosis in strongly-pigmented mugmel2 cells. Although there was a trend indicative of sensitization of SK-Mel28 cells to erastin by melanin depletion, this was presumably negated by the strong effects of erastin on the mel+ population.

Furthermore, we showed that there was a melanin-dependent sensitivity to ferroptosis induction, when melanin-depleted mugmel2 cells were compared with pigmented mugmel2 cells. Ferroptosis was induced either directly through the inhibition of GPX4 or indirectly by blocking the glutamate antiporter function of the system Xc- (erastin) and concordant perturbation of VDAC function. We found that mugmel2, derived from an NRAS-driven CM, is largely refractory to ferroptosis induction *in vitro* under conventional conditions, whereas these cells can be sensitized through depletion of glutamine (a co-factor for GPX4 function). The ferroptosis refractory nature of this RAS-driven cell line is contra-dogmatic, as both RSL3 (RAS specific lethal 3) and erastin have been selected through a RAS hyperactivation specific *in vitro* synthetic lethal screen^37^. Therefore, we reason that chemical depletion of melanin sensitizes mugmel2 cells to ferroptosis. This highlights the functional relationship between the presence of melanin and ferroptosis resistance, *in vitro*. Ferroptosis induction through GPX4 inhibition (with RSL3) or concordant inhibition of XCT and VDAC1/2 (with Erastin) generally lead to a reduction of metastasis formation, in both pigmented and non-pigmented melanoma cells. A significant difference between pigmented and non-pigmented mugmel2 cells, could be seen upon RSL3 treatment, but not upon treatment with erastin, indicating that non-pigmented cells are more susceptible to canonical ferroptosis induction through inhibition of GPX4. Conversely mugmel2 cells were strongly susceptible to ferroptosis induction with erastin independent of pigmentation.

These findings led us to search for both elevations in GPX4 levels or alterations in melanin-biosynthesis genes (*MITF, TYR, DCT, TYRP1*) in patient survival data, where we found that a high expression of GPX4 in CM significantly correlated with decreased overall survival. A high tumor expression of all melanin biosynthetic genes was correlated with increased death in CM patients. Strikingly, there were only insignificant reductions when correlating ferroptosis mediators *GPX4* and *SCL7A11* to overall survival of CM patients, on a whole tumor level. This however does not exclude the presence of inter-tumoral differences.

For UM we found that a higher GPX4 expression strongly correlated with shorter survival, whereas melanin biosynthetic activity when assessed as a measure of mRNA levels of TYRP1 showed an even stronger negative relation with survival of primary UM patients. No transcriptomics data was available to us for CoM melanoma at the time of writing.

VDAC1, one of the putative targets of erastin, showed a significant negative correlation relation with patient survival for both CM and UM. This finding is in line with the strong inhibitory effect of erastin and explains erastin’s strong inhibitory capacity on circulating cancer cells irrespective of intracellular melanin levels. Our findings are in line with Nawarak et al 2008, who show that arbutin, a known skin whitening agent and inhibitor of TYR, works by enhancing VDAC1 protein levels in A375 melanoma cells^38^.

Taken together, our findings establish a functional link between intracellular melanin levels in melanoma cells irrespective of their tissue of origin. Cells containing melanin survive longer in the circulation of zebrafish during experimental micro-metastasis formation and hence display an enhanced capacity to establish micro-metastatic colonies. In line with the elegant experiments performed by Pinner et al in 2017^22^, melanin content lowers (albeit in our hands not significantly so) the migration capacity of pigmented cells (endogenously or exogenously pigmented). Here our findings prove that metastatic dissemination and metastatic initiation is effectively enhanced by the presence of intracellular melanin.

The biological function of melanin as a ROS quencher is widely accepted^13,14,19^. Furthermore, there are several studies that correlate melanin concentrations, either through direct measurements of melanin levels or through the detection of blood borne mRNA in CM patients, with a worse prognosis^21,36^. Paradoxically, there are experimental studies showing a converse role of melanin, inhibiting small scale metastasis in animal models^21,22^. Taken together, we reason that our findings bridge the gap between the observed phenomenon in patients and the discrepancy in experimental animal models by showing that pigmented melanoma cells have a survival advantage *in vivo* in the blood circulation and are more resistant to ferroptosis. Furthermore, we show, using available patient survival data (TCGA and LUMC cohort for UM), that CM, UM patients have a worse prognosis when melanin biosynthesis is upregulated, notably the expression of terminal melanin biosynthetic enzyme TYRP1. VDAC1 was identified as another gene associated with a negative disease outcome and is a potential target for future therapy.

## Materials and methods

### Ethics statements

All animal experiments were approved by Animal Experiments Committee (Dier Experimenten Commissie, D.E.C.) under license AVD1060020172410. All animal were maintained in accordance with local guidelines using standard protocols (www.ZFIN.org)

### Stable cell culture

Cell line SK-Mel28 (CVCL_0526) was acquired from ATCC, while other cell lines were kindly provided by Dr. B. Rinner (Mugmel2, CVCL_JQ50)^31^, Prof. Dr. M.J. Jager (CRMM1 (CVCL_M593) and CRMM2(CVCL_M593))^43^, Dr. A.G Jochemsen (OMM2.3 (CVCL_C306))^44^ and Dr S. Alsafadi (XMM66, CVCL_4D17)^45^. Human CM PDX derived cell lines PDX11917 (alternatively named, M011-5.X1.CL) was kindly provided by Prof. D. Peeper^34^. All cells used were routinely imaged or observed using an inverted automated EVOS microscope (Thermo Scientific, Waltham, USA) using eGFP and RFP filters to ensure retention of normal phenotypes and to verify fluorescent tracer expression

Cells were cultured in a humidified incubator, 5% CO_2_ at 37°C, all cells were intermittently tested for the absence of mycoplasma using the universal mycoplasma detection Kit (American type cell culture (ATCC), LGC Standards GmbH, Wesel, Germany) according to the manufacturer’s prescriptions. All cells, with the exception of primary UM cells, were cultured in Dulbecco’s modified eagles’ medium (DMEM), enhancing melanin biosynthesis due to its high tyrosine levels (3,5-fold higher than RPMI1640). DMEM was supplemented with 10% fetal bovine serum and glutamax (GIBCO, Thermo scientific). Cells were propagated through subsequent medium removal, washing with Dulbecco’s phosphate buffered saline (DPBS) and incubation with 2mL tryplE (GIBCO). Cells were carefully dispersed after the addition of DMEM up to the original culture volume.

### Melanin measurement

Melanin was measured spectrophotometrically, after solubilization in 1M NaOH, containing 10% DMSO (v/v) as described by Friedman et al.^18^ In brief, cell pellets were collected of 2 × 10^6^ cells by tryplE incubation, inactivation and subsequent centrifugation. Cell pellets were stored frozen at -20°C prior to measurement. A standard curve of chemical eumelanin (Sigma, Zwijndrecht, the Netherlands) was made ranging from 1 mg/mL to 7,8125 µg/mL (2-fold dilution series) in triplicate. Standards and samples were solubilized by addition of 1M NaOH, 10% DMSO and incubation at 80°C for 30min. Melanin negative samples (breast cancer cells MDA-mb231 cells expressing tdTomato were used as a negative control) were taken along and were used to subtract backgrounds after measurement. Absorbance was measured at 420nm and plotted; concentration was inferred from the standard curve.

### Lentiviral over expression and shRNA construct generation

Lentiviral overexpression and shRNA constructs from the Sigma TRC mission library were kindly provided to us by Dr. M. Rabelink from the department of department of molecular cell biology, from the Leiden university medical center (Constructs detailed in supplementary table ST1). Lentiviral particles were generated as described previously by Heitzer er al 2019^33^. In brief Hek293T cells were grown to 80-90% confluency and transfected after a medium change with psPAX2, pMD2.G and the transfer plasmid of choice at a respective molar ratio of 1.3 pmol, 0.72 pmol, 1.64 pmol using 30uL lipoD293 on a 75cm2 culture flask. Cell culture medium was exchanged for 20mL complete DMEM 24 hours post transfection. Viral particles were harvested 72h after the original transfection.

### Chemical compounds and drugs

Erastin, RSL3, ferrostatin and mitomycin-C were purchased from Cayman chemical (Ann Arbor, Michigan, USA). PTU (1-Phenyl-2-thiourea) was purchased from Sigma (Sigma, Zwijndrecht, the Netherlands). BODIPY™ 581/591 C11 was purchased from Thermo scientific (Thermo Scientific, Breda, the Netherlands). Liperfluo was purchased from Dojindo (through GERBU Biotechnik, Heidelberg, Germany).

### Transmission electron microscopy sample preparation

Cells were cultured on thermanox (Thermo scientific/Nunc) coverslips, fixation was performed with a mixture of 2% glutaraldehyde and 2% formaldehyde in 0,1M Na-cacodylate buffer pH=7.2 Post-fixation was performed with 1% OsO_4_ +K_4_Fe(CN)_6_ (15µg/ml) in demineralized water for 1 hour at room temperature, after dehydration through a graded series of ethanol, all specimens were kept for 16 hours in epoxy resin (Agar Scientific,) before embedding. Ultrathin sections were collected on formvar-coated one hole copper grids. Electron microscopy images were obtained with a JEOL 1400Plus Transmission Electron Microscope (Tokyo, Japan) at 80KV.

### Chemical melanin depletion

Commonly PTU (Sigma) is used, dissolved in water, for the inhibition of melanation of zebrafish larvae. We reasoned that PTU could also be used to block the biosynthesis of melanin *in vitro*. To this end, we treated mugmel2 cells with a concentration range of 1 - 0.0625 mM, dissolved in dimethylsulfoxide (DMSO, Sigma).

### Co-culture experiments

For the melanin transfer co-culture experiments we cultured highly melanotic cell line mugmel2^31^ expressing eGFP in the absence and presence of phenylthiourea (PTU) a generic inhibitor of melanin biosynthesis.^20,47^ After two passages the PTU treated cells were considered melanin depleted, where depletion was validated through the spectrophotometric measurement previously described. The cells were treated with 100 µg/mL mitomycin-C (Sigma) in culture medium, under normal culture conditions, for three hours. The cells were washed with sterile PBS and DMEM subsequently prior to harvesting. UM cells were harvested as previously described. For both UM-tdTOM lines 1 × 10^6^ cells were seeded in a 75 cm^2^ culture flask mixed together with melanin depleted cells (mel^-^) and with highly pigmented cells (mel^+^), the cells were cultured for 4 days at normal culture conditions. Prior to harvesting the cells were checked for presence of remnant eGFP signal (donor cells) and tdTOM (acceptor cells) and for presumptive melanin transfer. The cells were washed extensively, prior to harvesting as previously described.

### WST1 proliferation assay

Mugmel2 cells (7,5 × 10^4^) were seeded in 100 µL in flat bottom 96 wells plates (Corning), combining both cells with prior chemical inhibition of melanin biosynthesis and vehicle control (DMSO) treated cells in the same plate, in triplicate. Cells were left to attach overnight and were subjected to ferroptosis induction using erastin and RSL3, compared to DMSO control for 3 days. Proliferation was measured based on

WST1 conversion, following the manufacturers description. Values were normalized and plotted with vehicle treated control set to 100% survival.

### Zebrafish engraftment

Primary cells were dispersed as described previously by Groenewoud et al^46^. In brief, cells were harvested from adherent cultures through tryplE addition, and subsequently concentrated by centrifugation. Cells were transferred to a 15 mL centrifuge tube and centrifuged for 5 minutes at 200 x g, followed by complete removal of all DPBS. The cell pellet was resuspended in 1 mL DPBS and subsequently counted. The cells were pelleted again at 200 x g whereafter the DPBS was removed after centrifugation. To completely remove all DPBS the cells were centrifuged for another minute at 200 x g, the cell pellet was resuspended to a final concentration of 250 × 10^6^ · mL^-1^ in 2% polyvinylpyrrolidone 40 (PVP_40_) in DPBS.

In brief, cells were injected into zebrafish larvae of either *casper* or (Tg*(fli:eGFP x casper*) zebrafish larvae at 48hpf into the duct of Cuvier (doC) also known as the embryonic common cardinal vein using a capillary glass needle.

### Drug treatment of engrafted zebrafish

Fish were bred and maintained until 48hpf, whereafter they were injected with approximately 300-400 cells per individual, through the doC allowing the cells to disseminate hematogenously within several hours after injection. One hour post injection possible dead larvae were removed from the injected pool and the injected individuals were divided over clean Petri dishes, with approximately 100-150 individuals per dish. Approximately 16hpi the injected larvae were screened using a stereo epi-fluorescent microscope, all the unwanted phenotypes (uninjected, malformed) were discarded. All larvae were randomly assigned to experimental groups in a 24 wells plate, with at least 6 wells containing 6 fish per well per condition. After plating the fish, approximately 16-18hpi the fish were treated with the appropriate level of inhibitor dissolved in DMSO and diluted to the final concentration in zebrafish water.

### Zebrafish xenograft data acquisition and analysis

For kinetic measurements of tumorigenicity engrafted individuals were imaged at 1,4- and 6-days post implantation using an epifluorescent stereo microscope. At the first time point the microscope settings (exposure time and gain) were set on the control group of each sample population, taking care that signal saturation was not attained to allow for signal increase due to cell growth. Each sample set was imaged using the same settings throughout the duration of the experiment. All images were analyze using a custom imageJ MACRO (Zenodo DOI: 10.5281/zenodo.4290225). Data was normalized to the vehicle control group of each experimental population; two biological replicates were combined with at least 20 individuals per biological replicates.

### Zebrafish data acquisition and statistical analysis

All zebrafish larval engraftments were performed in biological duplicate, unless otherwise stated. All groups were >20 individuals per biological repeat, unless otherwise stated. All individuals were randomized and entered into either control or experimental groups, all individuals were randomly selected and imaged using the same exposure setting using a stereo fluorescent microscope. Outliers were removed from all data sets using GraphPad Prism 8.0, (Q5) prior to normalization and combination of all biological replicates. Data was normalized to either control (drug treatment) or to day one (in the case of growth kinetics experiments). Statistical significance was tested with an ANOVA, for normally distributed data sets, otherwise a Kruskal-Wallis test was used. Error bars depict ±SEM.

### qPCR of zebrafish derived cells

Mugmel2-dTomato cells were engrafted as previously described. At 1-, 4- and 6- days post engraftment the tails of approximately 300 zebrafish larvae were amputated after euthanasia. The resected tissue was collected and incubated in 0.4 mg/ml Liberase TL (Roche) and Primocin (Invivogen) in PBS for 20 minutes under intermittent agitation (approximately every 5 minutes). Enzymatic digestion was inhibited, through the addition of up to 10% FCS, after a single cell suspension was generated. Cells were concentrated by centrifugation (5 minutes at 200 x g). The cell pellet was resuspended in 1ml DPBS and filtered using a 50µm cell strainer (Corning). mugmel2^dTomato+^ cells were isolated using a fluorescence activated cell sorter (FACS). Isolated cells were cultured in complete DMEM for 24 hours. Whole RNA was isolated using the Qiagen RNeasy kit following the manufacturer’s instructions. cDNA synthesis and subsequent qPCR analysis was performed as previously described. The primer sequences were previously described in Groenewoud et al 2023 (doi: https://doi.org/10.1101/2021.10.26.465874)

### qPCR analysis

Cells were harvested (1×10^6^) by centrifugation (200 x g for 5 min at 25°C. Whole RNA was isolated using the Qiagen RNeasy kit (Qiagen) according to the manufacturer’s description, treating the isolate on-column with RNase free DNAse for 15 minutes at room temperature. Total RNA yield was quantified using Nanodrop 2000 (Thermo scientific, Wilmington, USA) and 1 µg RNA was used to synthesize cDNA using the iSCRIPT cDNA kit (Biorad, Hercules, USA) according the manufacturers description.

Detection was performed using the iQ5 QPCR apparatus (Biorad), using IQ green super mix (Biorad), for 35 cycles. Primers were diluted in PCR grade nuclease free water (Gibco) at a concentration of 100 µM. All primers were tested for, and passed an efficiency test prior to use and were used at a final concentration of 10 pmol.

Glyceraldehyde-3-phosphate dehydrogenase (GAPDH) expression level was used as an internal reference for each experimental primer set. Transcript levels were corrected for loading to GAPDH expression and normalized using the ΔCT method. All samples were measured in at least 3 biological triplicates.

### In vivo ferroptosis measurement

To assess the in vivo increase of membrane lipid peroxidation we stained mugmel2^tdTom+^ cells prior to engraftment for 1 hour with 1µM Liperfluo (Dojindo) while in adherent culture (in low serum medium, 2,5% FCS). Subsequently the cells were washed three times with PBS and harvested with trypsin EDTA (as previously mentioned). Approximately 250-350 cells were engrafted per *Casper* zebrafish larva through the duct of Cuvier. 1 hour post engraftment the zebrafish were screened for presence of cells in the tail (CHT area), 24 hours post engraftment 5 randomly selected zebrafish larvae were embedded into low melting temperature agarose on a glass bottom confocal dish (Willco dish, Willco wells, Amsterdam, the Netherlands). Laser power and gain was set on the control (mel^+^) sample, measures were taken with Leica sp8 confocal microscope (Ex = 488nm Em = 500-550nm). Liperfluo intensity was measured within tdTom^+^ cells in the CHT, finally all measurements were normalized to mel^+^, at least 5 individuals were measured in three separate experiments.

### FACS experiments

Mugmel2 cells were cultured in complete DMEM with and without PTU (250 μM). After at least 3 passages, both melanin pro and deficient cells were seeded in 12 wells tissue culture plates. After 24 h, the culture medium was exchanged for complete culture medium containing Erastin (5 μM), RSL3 (10 μM), Erastin (5 μM) + Ferrostatin-1 (2 μM) or RSL3 (10 μM) + Ferrostatin (2 μM) complete culture-medium. After 8 hours of treatment, the medium was removed and cells were washed again with PBS. Bodipy C580/591, from the lipid peroxidation kit (Sigma) was added to the cells at a final concentration of 10 μM. Subsequently the cells were incubated for 30 minutes at 37°C. The media was removed and the cells were washed three times with PBS. The cells were collected by trypsinization and the shift in BODIPY C580/591 fluorescence intensity was measured at excitation/emission of 488/510 nm (FITC filter set). Data was analyzed with FlowJo 10.8.1.

### Patient data analysis

LUMC cohort: Genetic information on TYR, TYRP1 and DCT and information on the chromosome 3 status and BAP1 status was obtained from a database of 64 UMs in eyes enucleated at the Leiden University Medical Center between 1999 and 2008.

TCGA cohort: Information for both uveal and cutaneous melanoma patients were gathered from The Cancer Genome Atlas (TCGA), which is a publicly available database available at https://www.cancer.gov/tcga. The TCGA database for UM contains 80 patients and the TCGA database for cutaneous melanoma contains 458 patients. Data was accessed and analysed through GEPIA2^42^.

### Uveal melanoma patient samples

UM tissue was obtained from patients from the Leiden University Medical Center (LUMC) in Leiden, The Netherlands. Part of the tumor was snap frozen with 2-methyl butane and used for DNA and RNA isolation, while the remaining tumor tissue was fixed in 4% neutral-buffered formalin and embedded in paraffin.

For a gene expression array, material was obtained from 64 patients who underwent an enucleation for UM between 1999 and 2008, of which 51% were male and 49% female. The mean age at the time of enucleation was 61 years. The mean follow-up time (defined as the time period between enucleation and death) was 83 months (range 2 to 229 months). Follow-up was updated in 2020. At the end of follow up, 17 (27%) patients were alive, 37 (58%) patients had died because of metastasis, four (6%) had died because of other causes and six (9%) had died but the cause of death was unknown. Gene expression was determined with the Illumina HT12v4 array (Illumina, Inc., San Diego, CA, US). As published by de Lange et al 2015^39^.

Fresh tumor material was obtained directly after enucleation to establish spheroids. We also assessed mRNA levels of tumors included in the TCGA database (n=80) as published by Robertson et al 2017^40–42^.

### Institutional Review Board Statement

The analysis was approved by the METC of the LUMC (B14.003/SH/sh Approval Biobank OOG-2 “Oogtumoren (of een verdenking hierop)”, protocol Uveamelanoom-lab B20.026, approval June 2020). Fresh material was used for spheroids under METC protocol UM CURE 2020: Prospective collection: new treatment options for metastatic uveal melanoma (NL57166.058.16). The research adhered to Dutch law and the tenets of the Declaration of Helsinki (World Medical Association of Declaration 2013; ethical principles for medical research involving human subjects). Each patient had signed an informed consent.

### Patient data and statistical analysis

The statistical analyses were carried out with SPSS version 25 (IBM corp). Correlation between melanin-related genes and chromosome 3 status and BAP1 status were calculated with Mann-Whitney U test. The survival analysis was carried out using Kaplan Meier survival curves and splitting the gene expression data in the middle and comparing the 32 patients with lower TYR, lower TYRP1 and lower DCT with the 32 patients with higher TYR, higher TYRP1 and higher DCT respectively. In the LUMC cohort, survival was calculated with melanoma-related mortality as the endpoint.

The TCGA cohorts for both UM and cutaneous melanoma were analyzed with the interactive web server GEPIA2, splitting the population along the median for each gene. In these cohorts, survival was calculated with overall survival as the endpoint.

## Supporting information

Supplemtary figure 1. Transfer of melanin after co-culture

Supplemtary figure 2. TEM visualization of transferred melanin after co-culture

## Acknowledgements

We kindly thank Emilie Vinolo for the managerial assistance provided during this UMcure2020 project.

This work has received funding from the European Union’s Horizon 2020 research and innovation program under grant agreement No 667787 (UM Cure 2020 project, www.umcure2020.org).

We kindly thank Prof. R. Hoeben and M. Rabelink (Department of Cell Biology, LUMC) for providing shRNA (lentiviral) vectors (TRC library, Sigma-Aldrich)

We kindly thank Dr. Anne-Roos Schrader for her help in scoring the uveal and cutaneous melanoma stainings shown in figure 2

All graphics (excluding scientific data) were generated using Biorender.com

The results shown here are in whole or part based upon data generated by the TCGA Research Network: https://www.cancer.gov/tcga accessed through gepia(2)

AG conceived/designed, performed, analyzed all experiments (unless otherwise stated) and interpreted the data, wrote the manuscript.

MCG performed the UM patient pigment data analysis, read and provided feedback on the manuscript.

JY performed *in vitro* proliferation assays, read the manuscript.

GZ performed *ex vivo* qPCR measurements and ferroptosis FACS experiments and read the manuscript

GEML performed all the TEM sample prep and data acquisition, read the manuscript.

MJJ supplied materials and reviewed the manuscript.

FBE provided funding and reviewed the manuscript.

BES-J provided funding, supervised the project and reviewed the manuscript.

## Notes

### Competing Interest Statement

The authors have declared no competing interest.

