## Supplementary figures and images for "Melanin enhances metastatic melanoma colonization by inhibiting ferroptosis"

### Supplemtary figure 1. Transfer of melanin after co-culture

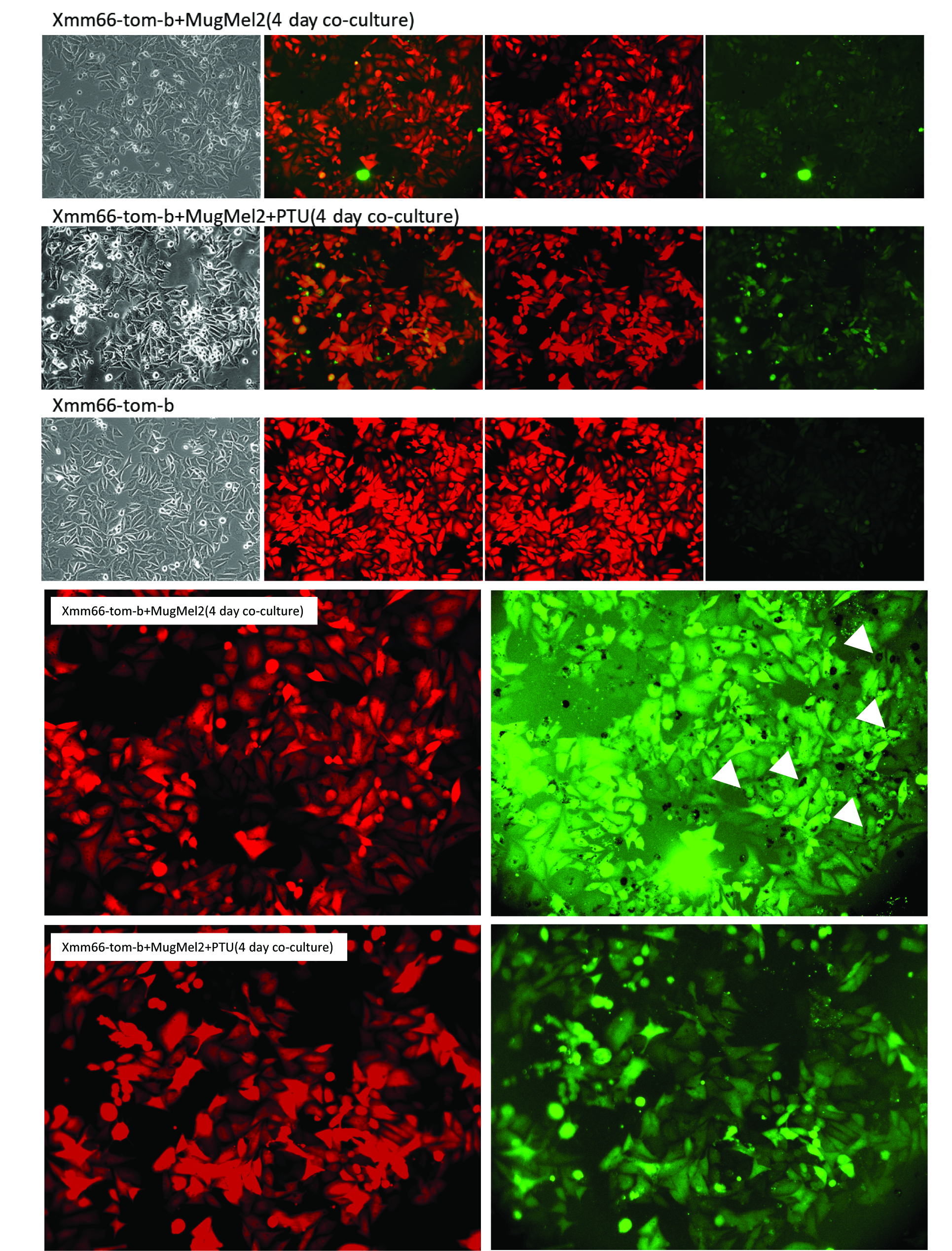

### Supplemtary figure 2. TEM visualization of transferred melanin after co-culture

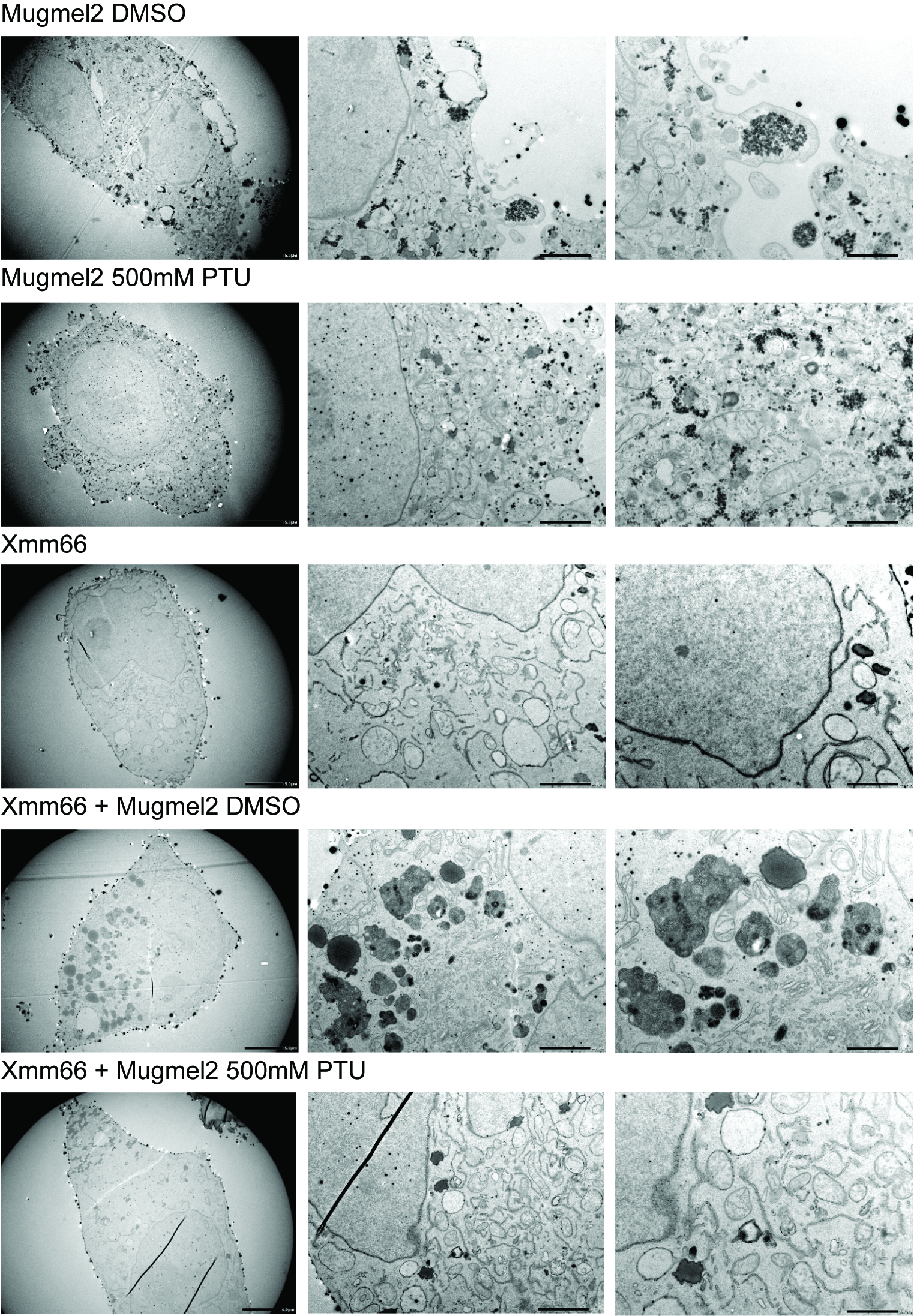
