## Supplementary table 1 for "Melanin enhances metastatic melanoma colonization by inhibiting ferroptosis"

| Plate | Well | Clone | Ref.Seq. | Gene Symbol | Organism | Validated | Knockdown | Gene Description |
| --- | --- | --- | --- | --- | --- | --- | --- | --- |
| AAK51 | B09 | TRCN0000083538 | NM_000372 | TYR | Human | not validated |  | tyrosinase  (oculocutaneous albinism IA) |
| AAK51 | B10 | TRCN0000083539 | NM_000372 | TYR | Human | not validated |  | tyrosinase  (oculocutaneous albinism IA) |
| AAM10 | C08 | TRCN0000097681 | NM_011661 | Tyr | Mouse | Yes | 0.76 | tyrosinase |
| AAM10 | C11 | TRCN0000097684 | NM_011661 | Tyr | Mouse | Yes | 0.87 | tyrosinase |
